# The canine hair metabolome encodes age and coat exposure in over 700 dogs

**DOI:** 10.64898/2026.11.10.765505

**Authors:** Fayzan Chaudhry, Lia Ficaro, Tori Rodrick, Bhupendra Singh, Aman Singh, Brenna Swafford, Drew Jones, Jan Krumsiek

## Abstract

Hair can be collected noninvasively and stored for years, making it a practical sample for studying physiology and environmental exposure. Yet the chemical information it carries in dogs remains largely unexplored. We profiled 782 banked hair samples from 700 Golden Retrievers by untargeted liquid chromatography–mass spectrometry, detecting 4,533 compound groups and revealing an unexpectedly rich chemical profile. Prominent signals reflected skin-surface and hair chemistry, including amino acids, peptides and urocanic acid, alongside surfactants, plasticizers and pesticides detectable despite washing. A signal annotated as imidacloprid distinguished dogs receiving topical products containing this insecticide from dogs receiving no flea or tick preventive (AUC 0.865). Hair chemistry also varied with age, most strongly between puppyhood and early adulthood. Oxidized lipids generally declined, whereas amino acids and peptides remained largely stable. A multivariate model estimated age with moderate accuracy (cross-validated r = 0.66) and assigned a higher age to the later sample in 64 of 75 repeatedly sampled dogs. Because age and collection date were strongly correlated, their effects could not be fully separated, although date-restricted analyses supported an age-related contribution. Together, these findings establish banked dog hair as a promising resource for studying chemical exposure and life-stage-associated changes in chemistry.

## Introduction

Companion dogs provide a natural model for studying aging. Their lifespans vary widely, veterinary medicine and canine genomics are well established, and they age alongside their owners in shared environments^1,2^. These shared environments also make dogs potential sentinels of chemical exposure. In one study, silicone tags worn by dogs and wristbands worn by their owners captured correlated profiles of household chemicals^3^. Studying aging and environmental exposure together across large canine cohorts, however, requires a biospecimen that is easy to collect and suitable for long-term storage.

Hair meets both requirements. It can be collected noninvasively, potentially by owners themselves, and stored frozen. Because hair grows slowly, a single clipping can retain chemical information accumulated over weeks to months, rather than provide the snapshot offered by blood; clipped hair in Labrador retrievers, for example, takes 14–15 weeks to regrow^4^. Hair cortisol is an established marker of longer-term physiological activity in humans^5^ and has also been validated in dogs^6,7^. Storage conditions nonetheless matter: cortisol, other steroids and lipid mediators decline in human hair kept at room temperature^8^, although cold storage may limit these losses.

Untargeted metabolomics extends hair analysis beyond individual compounds to broad chemical profiles^9^. In humans, an early study distinguished pregnancies complicated by fetal growth restriction from healthy pregnancies using hair metabolomics^10^. Subsequent liquid chromatography–mass spectrometry (LC-MS) studies have detected thousands of features reflecting endogenous metabolism and environmental exposure^11^, while lipidomic analyses have traced hair lipids to both sebum and blood ^12^. In rats, sebaceous lipids extracted from fur vary more with sex and age than with body region or among individual animals^13^, suggesting that fur chemistry may reflect age-related physiological changes.

In dogs, by contrast, hair analysis has largely focused on selected compounds, including cortisol^7,14^, nicotine^15^, bisphenol A^14^, persistent organic pollutants^16^ and per- and polyfluoroalkyl substances^17^. One validated method also screened for 394 pesticides, veterinary drugs and other residues^18^. Untargeted studies have been limited to volatile compounds measured by gas chromatography in cohorts of 36 dogs or fewer^19,20^. To our knowledge, no cohort-scale untargeted LC-MS study of canine hair has been reported, despite the availability of banked samples from large canine cohorts; the Dog Aging Project precision cohort alone has collected hair from 976 dogs^21^.

The Golden Retriever Lifetime Study provides an opportunity to address this gap. Between 2012 and 2015, it enrolled 3,044 Golden Retrievers in the United States, aged 6 months to 2 years, and has since collected hair and other biological samples at annual veterinary examinations, alongside clinical and owner-reported data^22,23^. We profiled 782 banked hair samples from 700 dogs by untargeted LC-MS in a single campaign, including repeated samples from 75 dogs collected years apart. We assessed measurement quality, characterized the chemical composition of canine hair, tested its ability to capture a recorded chemical exposure, and examined associations with age and their underlying chemical signatures. Because age and collection date are closely linked in this cohort, we also evaluated their potentially confounded effects.

## Results

### A cohort-scale hair metabolome in 700 dogs

We analyzed 782 banked hair samples from 700 Golden Retrievers in the Golden Retriever Lifetime Study (Figure 1A). The dogs were born between April 2010 and August 2014, and samples were collected between August 2012 and February 2025, mostly at annual veterinary examinations. Ages at sampling ranged from 0.5 to 13.1 years (median 5.9; Figure 1B), with similar contributions from females and males and a predominance of neutered dogs. Seventy-five dogs were sampled two or three times, 0.4–9.5 years apart (median 3.9). Samples had originally been selected for a nested case–control study of 15 health conditions (683 condition-recruited samples and 99 controls). Because recruitment groups differed in age (median 3.3–11.1 years), we accounted for this in subsequent analyses. Samples were shipped for analysis in February 2026, and LC-MS data were acquired between 31 March and 21 July 2026. Estimated biorepository storage durations, calculated to a common reference date of 1 February 2026, ranged from 1.0 to 13.5 years (median 7.2), with the youngest dogs’ samples stored longest.

**Figure 1.**
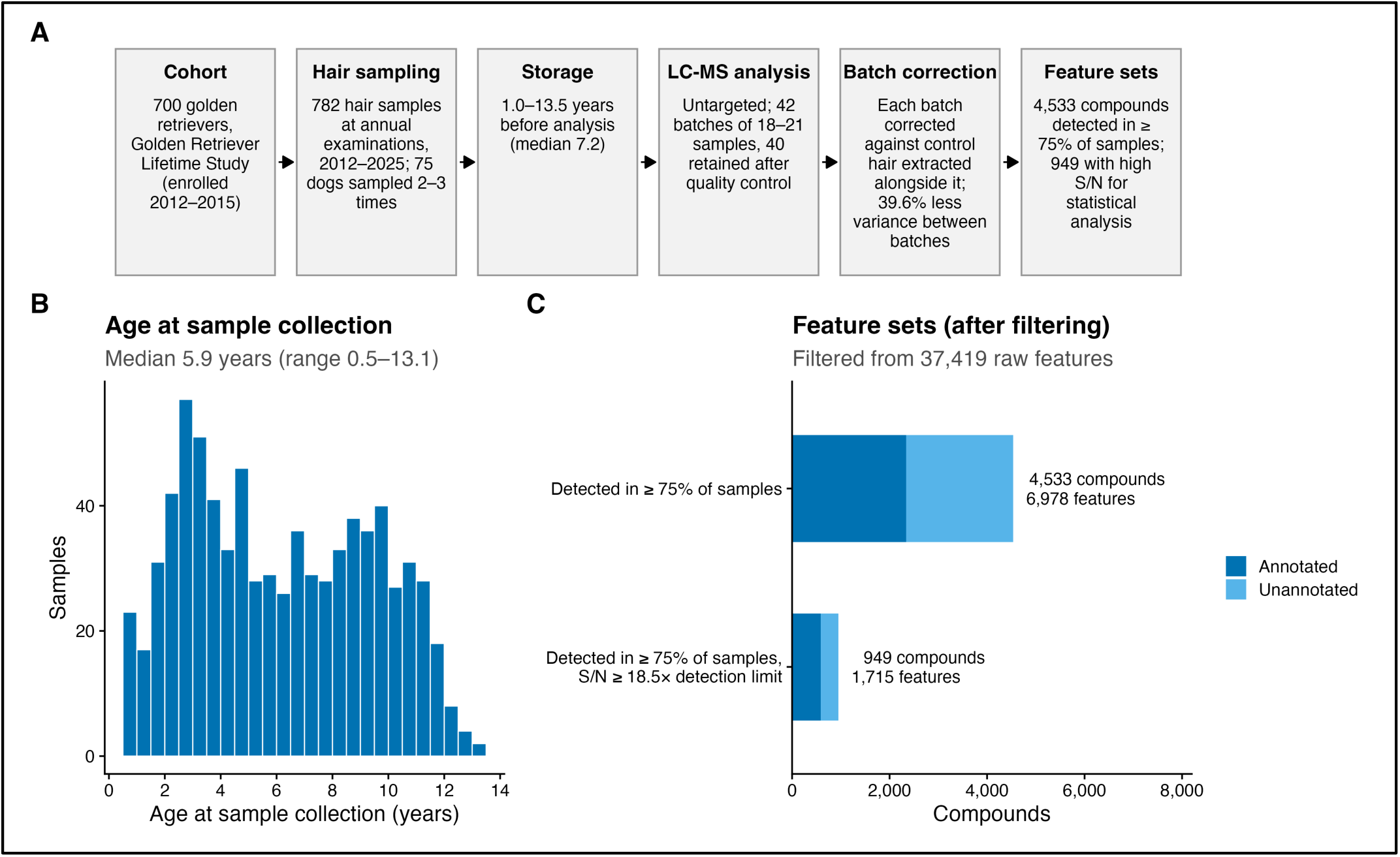
Study design. (A) Cohort, hair sampling, storage, LC-MS analysis, batch correction and feature sets. (B) Age at sample collection (median 5.9 years; range 0.5–13.1). (C) Compounds in the two feature sets: the catalog set (4,533 compounds, 2,343 annotated) and the analysis set (949 compounds, 589 annotated), with the number of features each comprises. A compound is a group of co-eluting related signals (main ion, isotopologues, adducts and alternative library matches). Feature selection is described in Methods.

LC-MS acquisition comprised 42 batches of 18–21 samples, each with blanks and two types of pooled-hair controls (Methods). Batch order was unrelated to age (r = 0.049). Quality control identified two batches in which the facility’s non-detection assignments disagreed substantially with raw intensities; these were excluded (Table S1), leaving 783 of the 822 initially measured samples in 40 batches (Table S2). One further sample had an unmatched identifier, giving the final dataset of 782 samples from 700 dogs.

The resulting hair profiles contained 4,533 putative compound groups detected in at least 75% of samples, of which 2,343 had tentative chemical annotations (Figure 1C; Table S3). We used this catalog set to characterize hair composition, and a more stringently filtered analysis set of 949 compounds, with stronger signals relative to the detection limit, to examine associations with age (Methods).

Batch correction required particular care because pooled hair can yield chemically heterogeneous control extracts. Rather than rely solely on the facility’s normalization to previously extracted controls, we used controls extracted alongside each batch and remeasured together under common instrument conditions (Methods). The correction reduced between-batch variance by 39.6% in held-out samples across 4,000 features. In the analysis set, the median share of variance attributable to batch fell from 0.255 to 0.190 (26%), improving 1,089 of 1,715 features and eliminating an apparent 85-fold batch shift in the highest-signal compound (Figure S1; Supplementary Note 1). Notably, despite these improvements, some differences between batches remained, accounting for 26.4% of the total variation in the measured chemical profiles (PERMANOVA; Supplementary Note 1).

### Dog hair carries skin-surface chemistry and records topical exposure

Most of the compounds detected in hair were shared across dogs. Each sample contained an average of 4,088 of the 4,533 compound groups, and the median compound was detected in 92% of samples (Figure 2A). Their contributions to the LC-MS signal, however, varied widely. Ranked by median peak area, the 10 compounds with the largest signals accounted for 30% of the total signal, and the top 100 for 66% (Figure 2B).

**Figure 2.**
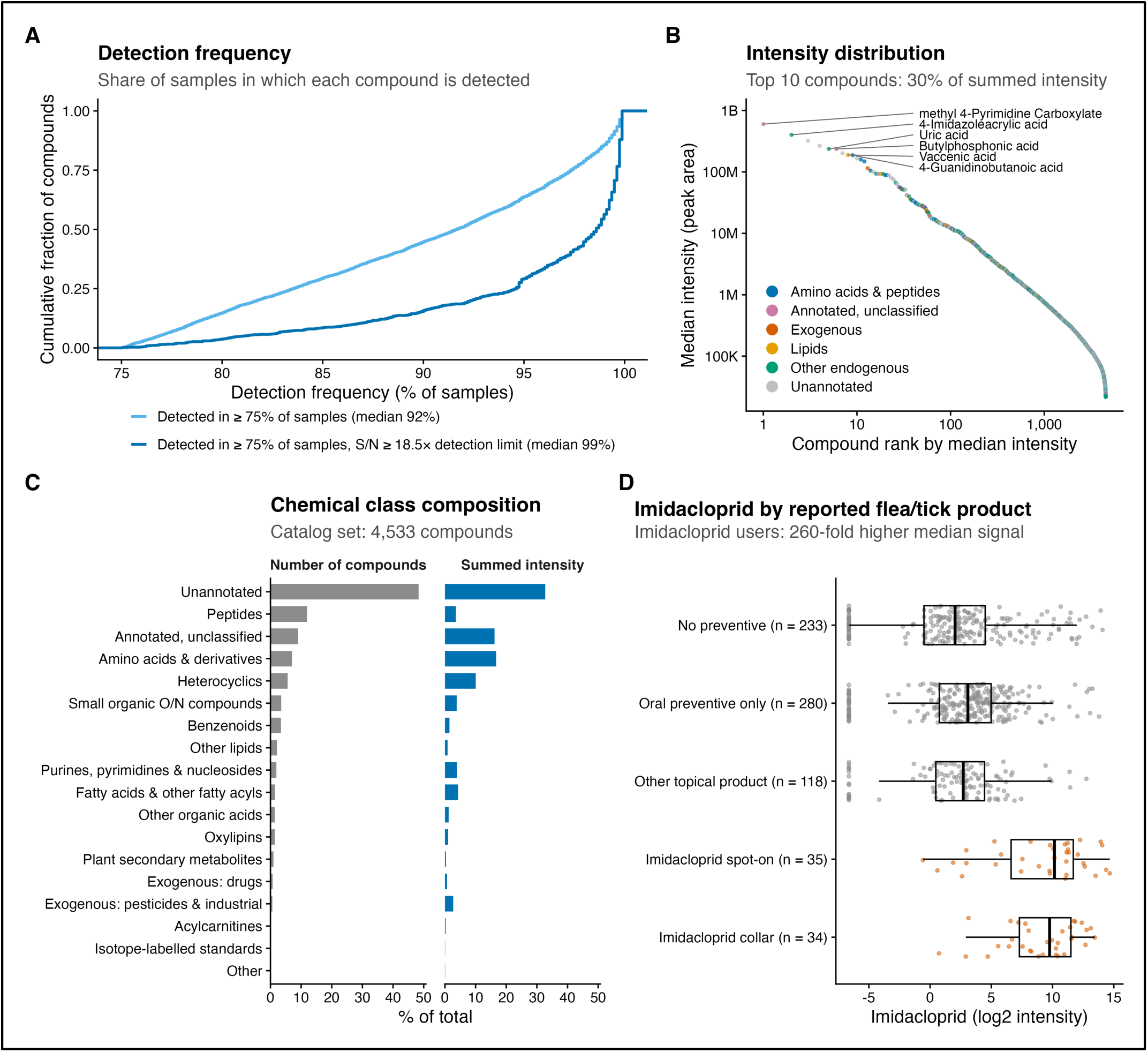
Composition of the dog hair metabolome. (A) Detection frequency of compounds in the catalog set (median 92%) and the analysis set (median 99%). (B) Compounds ranked by median raw signal; the 10 compounds with the highest signal carry 30% of the summed signal, and the six highest annotated compounds are labeled. (C) Chemical classes of the catalog set as a share of compounds and as a share of summed signal (the sum over compounds of each compound’s median raw peak area). (D) Signal of the feature annotated as imidacloprid in each dog, by owner-reported flea and tick product in the month of collection: no preventive (233 dogs), oral preventive only (280), other topical product (118), imidacloprid spot-on (35) and imidacloprid collar (34). Values at the far left are non-detects. Classes are from the HMDB chemical taxonomy and name rules (Methods); annotations are MS/MS library matches.

Much of this chemistry was characteristic of the skin surface and hair shaft (Figure 2C; Table S4). Among the 2,343 tentatively annotated compounds, peptides, mainly di- and tripeptides, were the largest class (542 compounds, 23%), consistent with the breakdown of hair proteins. Amino acids and their derivatives accounted for 316 compounds (13%) and contributed the largest share of measured signal (16.8%). Free amino acids, together with urocanic acid, pyroglutamic acid and lactate, are constituents of the skin’s natural moisturizing factor^24^; urocanic and pyroglutamic acids have also been measured on canine skin^25^. Heterocyclic compounds (253, 11%) contributed another 10.1% of the signal, largely through urocanic acid and an unidentified compound with the same molecular formula. Citrulline was consistent with proteins of the hair shaft^26,27^, while fatty acids, sphingoid bases and oxidized lipids were consistent with sebum. Purine metabolites, including uric and allantoic acids, were also prominent, consistent with dogs breaking purines down past uric acid to allantoin^28^. Individual compound identities remain tentative without authentic standards (Supplementary Note 2).

Not all of the detected chemistry originated from the dog itself. Despite extensive washing before extraction, hair contained surfactants, plasticizers, preservatives and pesticides. Four of the 30 annotated metabolites with the largest LC-MS signals were sulfate surfactants of the kind used in grooming products, and another was putatively annotated as Surfynol PSA-336 (Table S4). Other compounds included phthalates and bisphenol A, which leach from dog toys and training devices^29^, as well as methylparaben and several pesticides. We also detected compounds potentially derived from diet, including solanidine, an alkaloid found in potatoes whose serum levels track potato intake in people^30^ (Supplementary Note 3).

To test whether hair chemistry reflected a documented exposure, we examined imidacloprid, an active ingredient in topical flea and tick treatments. Selected before testing, the signal annotated as imidacloprid was substantially higher in dogs whose owners reported using an imidacloprid collar or spot-on product than in dogs receiving no preventive (69 versus 233 dogs; AUC 0.865; p = 2.9 × 10⁻²⁰; Figure 2D; Table S5). The result was similar for collars and spot-on products separately (AUC 0.878 and 0.852) and when compared against dogs receiving oral preventives (AUC 0.873; Table S6). By contrast, dogs receiving oral preventives or other topical products without imidacloprid showed little difference from untreated dogs (AUC 0.565 and 0.53). The association also persisted within calendar months (AUC 0.874; Table S7). Because imidacloprid can remain on the coat for weeks after treatment but is detected in blood for only days, the hair signal most likely reflects surface residue^31–33^. Detection in dogs without reported use may reflect incomplete records or transfer between animals^34^.

### Hair chemistry differs with age, especially early in life

We next examined whether hair chemistry varied with age. Because the samples came from groups recruited for different health conditions, which also differed in age (Figure 3A), we assessed associations both before and after accounting for recruitment group. Among the 949 compounds in the age-analysis set, 155 (16%) were associated with age at a false discovery rate of 5%: 92 had lower signals in older dogs and 63 had higher signals (Figure 3B). Individual associations were weak, with no absolute correlation exceeding 0.32 (Figure 3B; Table S8). After adjustment for recruitment group (i.e., disease status), 47 compounds remained associated, with effects about one-third smaller among compounds identified in both analyses. The overall pattern remained similar across alternative data normalizations and feature sets (r = 0.92–0.97; Table S9). Adjustments for batch, sex and measures of assay quality also gave broadly consistent results (Supplementary Note 4).

**Figure 3.**
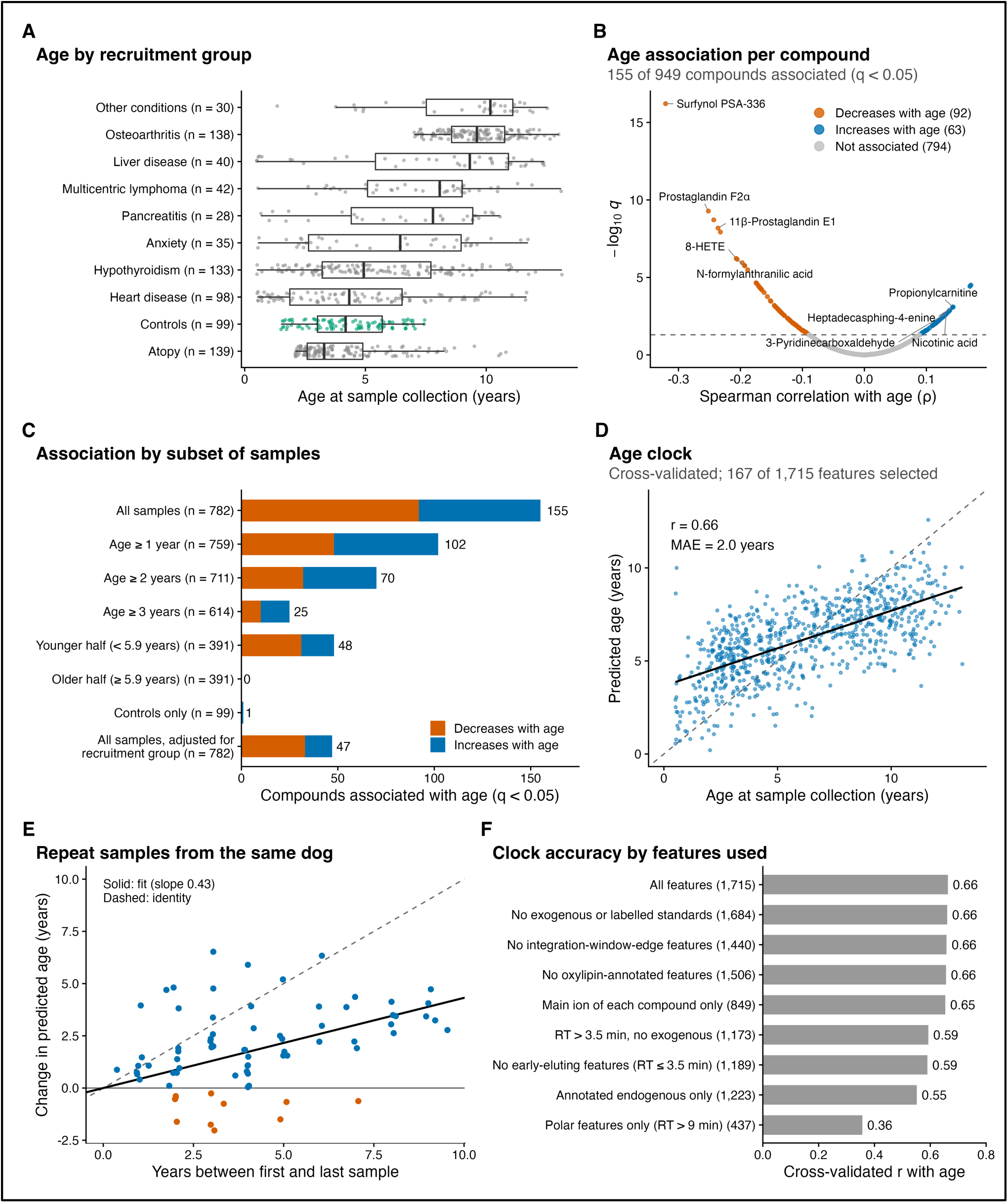
Association of hair chemistry with age. (A) Age by recruitment group (groups with at least 25 samples; the others pooled), with control dogs highlighted. (B) Association of each compound with age, represented by its most age-associated feature: Spearman ρ against −log10 q; 92 compounds fall and 63 rise with age (q < 0.05), and 794 are not associated. (C) Number of compounds associated with age in subsets of samples (all; aged at least 1, 2 or 3 years; younger and older half; controls only) and after adjustment for recruitment group, split into those that decrease and those that increase with age. (D) Age estimated by the elastic-net clock in cross-validation against actual age (r 0.66, mean absolute error 2.0 years; 167 of 1,715 features retained). (E) Change in estimated age between the first and last sample of each of the 75 dogs sampled more than once, against the years between them (later sample estimated older in 64 of 75; slope 0.43). (F) Clock accuracy by the features used, with the number of features in parentheses.

The association with age was strongest early in life (Figure 3C). When dogs younger than 1, 2 or 3 years were excluded, the number of associated compounds fell to 102, 70 and 25, respectively. Forty-eight compounds were associated with age in the younger half of the cohort (under 5.9 years), but none in the older half. Together, these findings suggest that hair chemistry differs most between puppyhood and early adulthood, with relatively little additional age-associated variation detectable later in life.

Although individual compounds were only weakly associated with age, their combined signals were considerably more informative. We trained an elastic-net model using all 1,715 LC-MS features, evaluating its predictions by 10-fold cross-validation with folds grouped by dog, so that repeat samples from the same dog never appeared in both training and test sets. Estimated age correlated with actual age at r = 0.66, with a mean absolute error of 1.99 years (Figure 3D). The model selected 167 features (Table S10), and its predictions were compressed toward the cohort mean, overestimating the youngest dogs and underestimating the oldest. This compression is usual for biological age estimators; epigenetic clocks underestimate age in older people in the same way ^35^. Accuracy was unchanged when the model used a single representative from each group of strongly correlated features (absolute Pearson correlation ≥ 0.85; 746 groups; r = 0.66), showing that it was not inflated by one compound contributing several related signals. Repeated samples from the same dogs tested whether the model orders samples within an individual, not only across dogs. In 64 of the 75 dogs sampled more than once, the model estimated the later sample to be older (85%; p = 3 × 10⁻¹⁰; Figure 3E).

We refitted the model on restricted sets of samples and features to test whether its accuracy rested on a few dogs or on signals that cannot reflect dog biology (Tables S11 and S12). Excluding dogs under one year changed little (r = 0.637 against 0.647 on the same samples), as did removing features of external origin, isotope-labeled standards or oxylipins, using only the main ion of each compound, or removing features at the edge of their integration window (r = 0.654–0.661 against a baseline of 0.663; Figure 3F). Accuracy fell when the model was restricted by annotation status or chromatographic region: r = 0.552 using only annotated endogenous features, 0.590 without the early-eluting lipid region and 0.357 using only late-eluting polar features. Within the older half of the cohort it reached only r = 0.114. The age signal is therefore spread across many compounds, weighted toward early-eluting lipids, and partly carried by unannotated features.

### Age-associated differences are concentrated in oxidized lipids

To identify the chemical changes underlying the association with age, we compared patterns across metabolite classes. Oxylipins, oxidized derivatives of polyunsaturated fatty acids and a known constituent of skin surface lipid^36^, showed the strongest and most consistent relationship with age. Their average standardized signal declined in older dogs (ρ = −0.19; Figure 4A). Of the 22 oxylipin compounds in the analysis set, 14 were significantly associated with age, and all 14 showed lower signals in older dogs (Figure 4B). These represented 10 distinct chromatographic peaks, as multiple compound annotations sometimes corresponded to the same measured signal ^37^. Unoxidized fatty acids showed a similar but weaker decline (class-level ρ = −0.08; linoleic acid ρ = −0.14), whereas amino acids and peptides showed no comparable trend (ρ = +0.03 and +0.04; Figure 4A).

**Figure 4.**
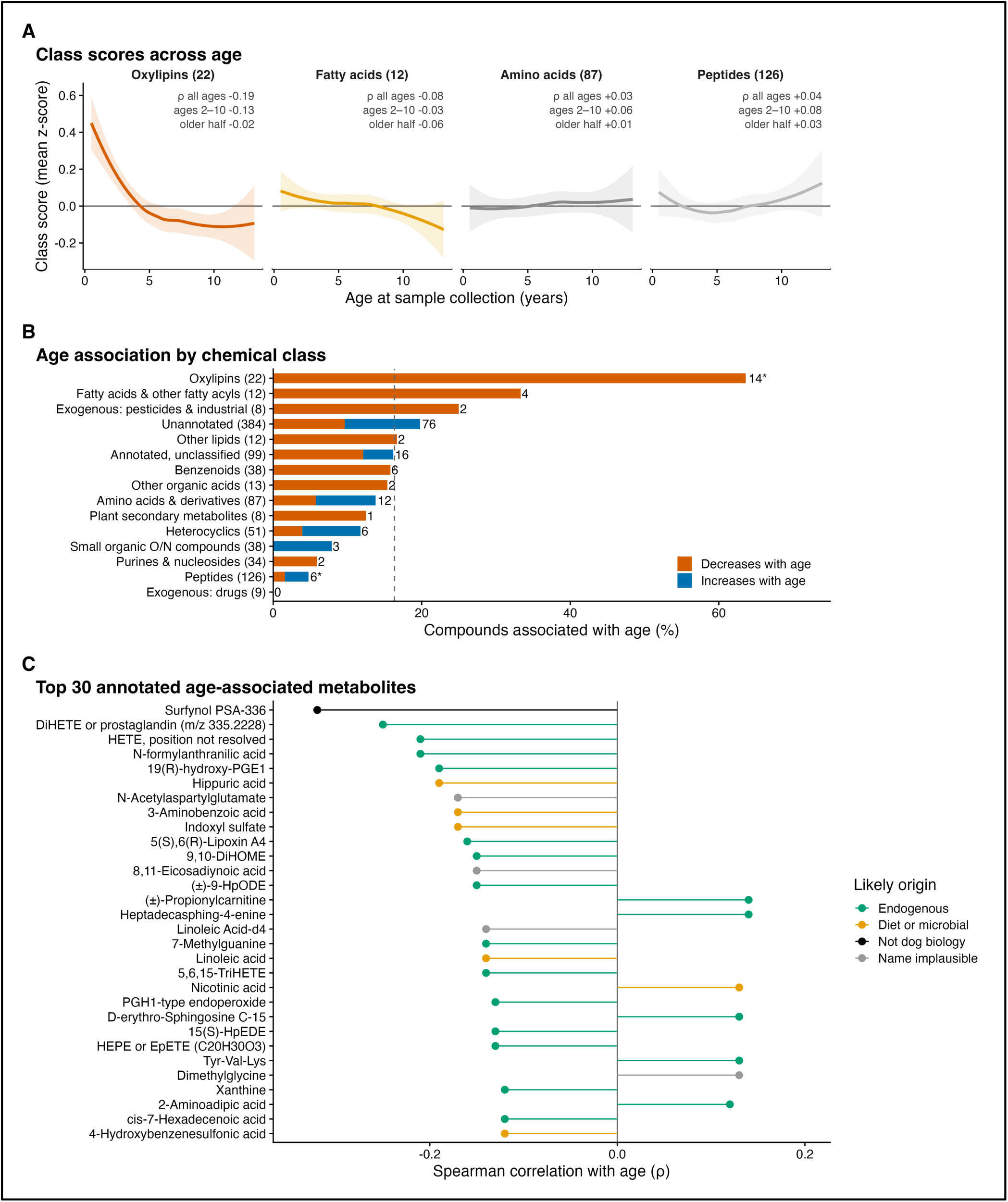
Chemical classes associated with age. (A) Class scores (the mean z-score of a class’s compounds) against age, shown separately for oxylipins, fatty acids, amino acids and peptides, with LOESS fits and 95% bands; the number of compounds is given above each panel, and within it Spearman ρ with age for all samples, for adults aged 2–10 years and for the older half of the cohort. (B) Share of compounds associated with age in each chemical class with at least 8 compounds, split into those that decrease and those that increase with age; the dashed line marks the overall 16%, and asterisks * mark classes that differ from all other compounds (q < 0.05). (C) The 30 metabolites of Table S13 by Spearman ρ, colored by likely origin. Names are putative (MS/MS library matches).

Several oxylipin signals were among the strongest individual age associations in the study (Figure 4C; Table S13). These included peaks tentatively identified as a prostaglandin or DiHETE (ρ = −0.25), a HETE (−0.21), 19(R)-hydroxy-PGE1 (−0.19), lipoxin A4 (−0.16) and 9,10-DiHOME (−0.15). Together, these annotations point to multiple pathways of fatty acid oxidation, including the cyclooxygenase, lipoxygenase and cytochrome P450 pathways. The first two are active in human sebaceous gland cells^38^, and skin microbes convert sebaceous fatty acids into oxylipins, among them the cytochrome P450 product 9,10-DiHOME^39^. Their precise identities remain uncertain, however, because structurally related lipids can produce overlapping signals. The decline was consistent with the broader age-associated pattern: it remained detectable among dogs aged 2–10 years (ρ = −0.13) but was nearly absent in the older half of the cohort (ρ = −0.02; Figure 4A).

Age-associated changes outside the lipid classes were less readily interpretable. Of the 155 compounds associated with age, 76 lacked chemical annotations. Among the annotated compounds, signals assigned to hippuric acid (ρ = −0.19), indoxyl sulfate (−0.17) and N-formylanthranilic acid (−0.21) also declined. These compounds may reflect dietary or gut microbial metabolism ^40,41^, but their signals co-eluted and varied together, limiting their independent interpretation (Supplementary Note 5). Other associations appeared to reflect changes in external exposure rather than the dog’s physiology. The strongest individual age correlation, for a compound annotated as the surfactant Surfynol PSA-336 (ρ = −0.32), disappeared after adjustment for collection date (ρ = +0.03). Age and collection date were strongly correlated (r = 0.954), but a date-restricted permutation test supported a contribution from age (P = 0.021; Figure S2, Supplementary Note 4).

The oxylipin findings were more robust to technical filtering: 12 of the 14 age-associated compounds remained significant after excluding features near the edges of their chromatographic integration windows. Taken together, these results identify oxidized lipids as the clearest chemical signature of age-associated variation in canine hair. Whether their decline primarily reflects changes in skin physiology during maturation, differences in exposure, or a combination of both remains unresolved.

## Discussion

Untargeted LC-MS profiling of 782 banked hair samples from 700 Golden Retrievers revealed a diverse chemical profile containing compounds associated with hair, skin and the external environment. The profiles captured a documented topical exposure and showed associations with chronological age, particularly early in life. These findings demonstrate the feasibility of cohort-scale canine hair metabolomics and its potential for studying chemical exposure and age-associated changes.

Much of the detected chemistry was consistent with the composition of hair and the skin surface, including amino acids, peptides, urocanic acid and lipids ^12,24,25^. Hair also contained surfactants, plasticizers and pesticides despite washing before extraction. These chemicals may be deposited directly on the coat rather than incorporated through systemic circulation. This is likely the case for imidacloprid, which persists on the coat after topical application^31,32^. The signal annotated as imidacloprid distinguished dogs receiving products containing this insecticide from dogs receiving no flea or tick preventive (AUC 0.865). This provides evidence that hair metabolomics can detect documented topical exposure, although its ability to measure other types of environmental exposure remains to be established.

Hair chemistry also varied with age, with the strongest differences occurring between puppyhood and early adulthood. Oxidized lipids showed the most consistent pattern, generally declining with age, whereas amino acids and peptides remained relatively stable. Oxylipins are present in skin-surface lipids and can originate from both host and microbial metabolism ^36,38,39^. Changes in skin and sebaceous activity during maturation could therefore contribute to these findings, although the underlying processes remain uncertain. Combining chemical signals allowed chronological age to be predicted with moderate accuracy (cross-validated r = 0.66), but prediction was poor among older dogs. The observed associations may therefore reflect changes during maturation more strongly than processes of late-life aging.

An important limitation is the strong correlation between age and sample collection date (r = 0.954). Because the dogs were born within a relatively narrow period, changes in grooming products, preventive use or collection procedures over the study period could resemble age effects. For example, the strongest individual age-associated signal was annotated as a surfactant, and its association disappeared after adjustment for collection date. A permutation analysis restricted to samples collected at similar times supported an age-related contribution, but the limited variation in age independent of collection date prevented the two effects from being separated reliably (Figure S2). Similarly, the increase in predicted age between repeated samples from the same dogs cannot distinguish aging from changes associated with collection date. Storage duration also varied systematically with collection date. Because the extracted chemistry is not expected to change once the hair is frozen, the more plausible calendar-time effect is a difference in what reached the coat before collection rather than degradation during storage.

Other limitations concern cohort selection and measurement. Samples were selected for studies of different health conditions, and adjustment for recruitment group reduced the number of compounds associated with age. The smaller control group did not independently establish the same associations. Residual batch effects, the restriction to one breed and uncertainty in compound identification further limit interpretation. In particular, the oxylipin annotations require confirmation with authentic standards, and the measured signals do not provide absolute concentrations.

Future studies should examine dogs of different ages sampled during the same period to distinguish age-related changes from calendar-time effects. Standardized collection protocols, targeted quantitative assays and confirmation of compound identities would strengthen the biological interpretation. Replication across breeds and prospective studies of health outcomes are also needed to determine whether hair chemistry provides information beyond chronological age. Overall, this study establishes banked canine hair as a useful resource for broad chemical profiling and detecting topical exposure, while identifying age-associated lipid changes that warrant further investigation.

## Methods

### Samples and storage

Hair came from the Golden Retriever Lifetime Study, a prospective cohort of 3,044 Golden Retrievers in the United States, enrolled at 6 months to 2 years of age between June 2012 and April 2015 ^22,23^. Each dog has an annual veterinary examination at which biological samples, hair among them, are collected and banked^23^. All hair samples analyzed here were collected with the study’s annual sample kits. The kit instructions ask for a lock of hair about 6 mm (¼ inch) in diameter and at least 5 cm (2 inches) long, cut as close to the root as possible from a site chosen by the owner, and placed in a screw-cap biorepository tube. Samples were cut by one of more than 3,400 participating study veterinarians or by the owner shortly before the visit; body site, collector and time from collection to freezing were not recorded, and this interval may have included time at room temperature or under refrigeration and overnight shipment to the biorepository. Once banked, hair was stored at −20 °C in cryogenic tubes, with no recorded freeze–thaw events or periods at ambient temperature, including during a transfer of the collection to a new biorepository in February 2024. Samples were shipped for analysis in February 2026.

The samples were chosen for a nested case–control study of 15 health conditions. Of the 782 samples analyzed, 683 came from dogs recruited under one of these conditions and 99 from control dogs. Eight conditions contributed at least 25 samples each (653 in total) and the other seven 1 to 10 each (30 in total). This study does not compare the health conditions with one another; recruitment group is used only as a covariate in the age analyses. Of the samples, 770 were collected at annual examinations, 4 at visits to repeat a sample collection, 5 at biopsies and 3 at necropsies. Of the 75 dogs sampled more than once, 68 were sampled twice and 7 three times (157 samples). Age at collection is the time from birth to the visit date, in years. The study records the visit date as the date of sample collection, although hair cut by the owner may predate the visit slightly. Birth dates were reported by the owners; enrollment required a three-generation pedigree^23^. All samples were extracted and analyzed in one campaign in 2026, with raw LC-MS data acquired between 31 March and 21 July 2026. We calculated approximate biorepository storage duration as the interval from collection to 1 February 2026, a nominal reference date at the start of the month in which samples were shipped for analysis. These estimates describe storage before shipment rather than the interval to extraction or LC-MS acquisition.

### Extraction and LC-MS

Hair was extracted and analyzed by untargeted LC-MS at the Metabolomics Core Resource Laboratory, NYU Langone Health (New York, NY). Extraction followed a protocol adapted from Chen et al.^11^. Each sample was washed two times for 1 min with 1 mL water:methanol (1:1, v/v), and the washes were discarded. The hair was dried in a vacuum concentrator (16 h, 45 °C), cut into 1-cm pieces and pulverized with three stainless-steel beads in a bead mill (BeadBlaster-24, 10-cycles 30s on, 30s off). The powder was extracted with 80% methanol in high purity water at a ratio of 30mg fur/mL buffer and containing a mix of doubly-labeled amino acids at a concentration of 1uM (catalog no. MSK-A2-1.2, Cambridge Isotope Laboratories). The powderized fur, beads, and buffer was transferred to a fresh beadblaster tube and homogenized for another 10 cycles, followed by centrifugation 21,000xg for 3 min. A fixed volume of supernatant (450uL) was dried in a speedvacuum concentrator and reconstituted in 100 µL methanol:water (1:1, v/v) followed by a bath sonication step (2min). The sample was again centrifuged (21,000 x g, 6 min) and 60 uL of the supernatant was transferred to glass LCMS inserts for data acquisition. These steps shape what is detected: in human hair, pulverizing yielded more metabolites than cutting, and the choice of extraction solvent changed the yield of most metabolites^11^.

Extracts were analyzed with fast polarity switching between positive and negative heated electrospray ionization. Briefly, all LC runs were carried out on a Dionex UltiMate 3000 system. The column was a Waters Atlantis Premier BEH Z-HILIC column (1.7 µm, 2.1 mm × 150 mm), with mobile phases A and B being 10 mM ammonium carbonate in water, LC-MS grade (A) and neat acetonitrile, LC-MS grade (B), respectively. The injection volume was 2 µL for all analyses. The flow rate was 0.1 mL/min, and the gradient was linear from 80% B to 20% B over 30 min, followed by a rapid return to 80% B over 1 min and equilibration at 80% B until 42 min. Total MS acquisition time was 30 min. The column oven was maintained at 25 ± 5 °C. The MS method was a Top5 polarity-switching method on a Q Exactive HF instrument, and the following settings applied to both polarities. For each cycle, a survey scan was carried out at 120,000 resolution with an AGC target of 3e6, maximum IT of 100 ms, and a scan range of m/z 67–1,000. For the data-dependent MS2 scans, the resolution was 15,000, the AGC target was 1e5, the maximum IT was 50 ms, the loop count was 5, the quadrupole isolation range was 0.4 m/z with an offset of 0.1 m/z, a fixed first mass of m/z 50, and an equal time stepped collision energy of 10, 30 and 80 nCE. Data-dependent settings were a minimum AGC of 1.0e4 with an intensity threshold of 2.0e5, apex trigger of 1–20 s, auto dynamic exclusion, and no charge, peptide or isotope exclusion. The source was heated electrospray ionization with a sheath gas flow rate of 35, aux gas flow rate of 15 and sweep gas flow rate of 0. Spray voltage was 3.5 kV for both polarities, with a capillary temperature of 320 °C and an aux gas heater temperature of 30 °C.

### Data processing

Untargeted metabolite features were detected with an in-house script “Ungrid”, resulting in tens of thousands of raw features. These features were filtered for minimum signal intensity (1e5), a unique m/z at a given retention time of 20 ppm, a 2 min retention window, and a minimum signal-to-noise ratio of 10 (Min_range). All candidate untargeted features were processed on a batch level, then the resulting features were consolidated across all batches based on their percentile ranking of the founding feature, equivalence by mzrt, RT_tol of 2.0 min, and 20ppm parent mass tolerance. The resulting deduplicated metabolite feature library was then relatively quantified in each of the batches independently to ensure feature alignment for inter-batch variation. The feature detection limit was defined per batch from the replicate analytical blank controls (Run design and quality control). Peak intensities were then normalized to the batch control replicates (per batch) to generate a ratio value with the sample as the numerator and the batch control average as the denominator.

Similarly, for putative identification of metabolites, all sample MS2 spectra were exported to .mgf format (per batch), and the spectra were identified by spectral library search against the NIST17MS/MS and METLIN tandem mass spectral libraries of metabolites using NIST MSPepsearchx64, along with their decoy libraries to determine the name FDR^42^. Search parameters were; `d a v l P /Z 0.01 /M 0.05 /MatchPolarity /HITS 1 /MinMF 1 /OutPrecursorType /OutIK /OutChemForm /OutPrecursorMZ /OnlyFound`. The top-scoring structure match for each data-dependent spectrum was returned using an in-house script wrapper “Dewey” for MSPepSearch_x64. Putative structures were sorted from high to low by their NIST score, and duplicate inchicores were discarded, retaining only the top-scoring MS2 spectrum and the neutral chemical formula and detected polarity (+ or −) of the putative metabolite. These chemical formulas were used to quantify the extracted ion chromatograms using the theoretical m/z of the library-annotated ion type (from the matching spectrum), 15 ppm, with an RT tolerance of 0.2 min using an in-house script (skeleton). Relative quantification of the peaks was carried out for each sample according to these criteria in a batch-wise fashion. The detection limit was defined the same as for untargeted features above, and all putatively named metabolites were normalized to the batch control replicates.

Putatively named metabolites and untargeted features were deduplicated and clustered using mz tolerance (10 ppm), RT tolerance (0.1 min), and Min_r2 (0.9) across the study samples. Redundant correlated features were discarded, while identifiable mass shifts (either named or un-named) were kept and assigned into clusters based on a standard table of common isotopes and adducts. Each resulting feature, named or untargeted, is thus the signal integrated within 15 ppm of one m/z and within a fixed retention-time window (0.5 min wide for 98% of features): at the theoretical m/z of the library-annotated ion for putatively named metabolites, and at the observed m/z for untargeted features. The facility delivered this processed table (33,072 features) and, separately, the same acquisitions before blank masking, clustering and normalization (37,419 features), which are the raw intensities referred to in these Methods. All scripts available on GitHub (https://github.com/NYUMetabolomics/plz).

### Run design and quality control

Study samples were analyzed in 42 batches of 18–21 samples. Control dogs were spread evenly across batches (2 or 3 per batch), and in the 40 retained batches, batch was unrelated to age, collection date, sex and recruitment group (Kruskal–Wallis and permutation χ² tests, all p ≥ 0.10). Each batch also included 8 blank injections (10 in two batches), 4 batch control injections and 4 day control injections (5 of each in two batches), following the usual design for untargeted LC-MS studies ^43,44^. Sample injection order was randomized using a custom script, with control blocks interspersed at fixed intervals throughout the acquisition and system suitability/conditioning injections before and after the study samples (n∼10). Two further controls were prepared from one shared pool of dog hair. Batch controls were extracted from the pool before the study and stored as extracts (n=150 parallel aliquots); day controls were pre-aliquoted as fur, but extracted together with each batch’s samples on the same day. Isotope-labeled amino acids served as internal standards. Blanks contained no hair and went through the same workflow side-by-side with all study samples and controls. After the last batch, the stored day-control extract of every batch was injected twice in one further run (84 injections), so each extract was measured once under the instrument conditions of its own batch and twice under common conditions.

For each feature in each batch, the detection limit was 3× the mean intensity of the batch’s blank injections + 10,000 intensity units; values below it were treated as not detected. For each batch, the facility’s marking of non-detected values was compared with this limit applied to the raw intensities; the two batches in which they disagreed were excluded (Results; Table S1), leaving 783 of 822 study samples in 40 batches. The retained batches were also ranked on deviation in detection rate, relative log-abundance offset and three measures of distance from the other batches (Table S2); none was excluded on this basis. One further sample was left out because its identifier did not match the cohort’s records. Batch membership was taken from the facility’s injection record.

### Batch correction

The facility normalized each feature in each batch to the mean of that batch’s batch control injections, a common form of correction based on quality-control samples^45^. Hair does not mix as a fluid does, so control extracts made from the same pool can differ in composition, and dividing by them can add differences between batches as well as remove them. We therefore anchored each batch to its own day controls and used the additional run to separate the instrument conditions of each batch from the composition of its day-control extract. The correction works on log2 raw intensities, one feature at a time, and only control samples set its direction.

Methods such as ComBat^46^ estimate batch-specific shifts in means and variances from the study samples themselves. When the composition of samples differs across batches, these estimates can conflate technical variation with genuine biological differences. We instead used the day controls to determine the direction of each batch correction independently of the study samples. The study samples were used only to estimate a single correction strength per feature, shared across all 40 batches, rather than separate adjustments for each batch. Neither age nor other characteristics of the dogs entered this estimation. We assessed the effectiveness of the correction on samples held out from the estimation of its strength.

For feature f and batch b, let d_fb_ be the mean log2 intensity of the batch’s day control injections minus the mean of these values over the 40 batches, and let p_fb_ be the mean of the two re-run injections of that batch’s day-control extract, centered across the 40 extracts. The additional run measured all extracts under common conditions, so p_fb_ reflects what is particular to the extract (its portion of hair and its extraction), and Δ_fb_ = d_fb_ − p_fb_ reflects the instrument conditions of batch b. Study samples need not respond to these conditions as the controls do, so each feature received one strength, s_f_: the least-squares slope through the origin of the study samples’ batch means (centered across batches) on Δ_f_, estimated from at least 10 batches and limited to the range 0–2. The corrected log2 intensity of a study sample in batch b is its log2 intensity minus s_f_ Δ_fb_. The strength depends on batch membership only, never on age or any other characteristic of the dogs, and was estimated once from all study samples. Because the extraction of each day control cancels along with its composition, the correction addresses instrument conditions, not differences in extraction between batches.

A feature was corrected this way if it was detected in the day controls of at least 38 of the 40 batches and in the additional run; for its missing batches, the deviation was replaced by that batch’s median deviation over all such features. The remaining features (357 of the 1,715 in the analysis set) kept the facility’s batch control normalization. To measure the effect of the correction without fitting and judging on the same samples, study samples were split in half within each batch; strengths were estimated from one half and the share of variance between batches assessed in the other (Results). An uncorrected matrix was rebuilt from the raw intensities for comparison (Figure S1).

### Feature selection and compounds

Features were selected on the raw intensities of the 40 retained batches, without reference to age or any other characteristic of the dogs. The first filter kept features detected in at least 75% of study samples, 8,688 of the 37,419 raw features. The second additionally required a median signal-to-noise ratio of at least 18.5 among the samples in which the feature was detected, where signal-to-noise is intensity divided by the detection limit; the threshold is the upper quartile among features passing the first filter, and 2,172 features passed both. Both sets were then restricted to features present in the facility’s processed table, giving the catalog set (6,978 features) used to describe composition and the analysis set (1,715 features) used for all statistics. Values are analyzed on the log2 scale, with non-detects set to the feature’s minimum.

Features assigned to the same putative compound share a compound identifier from the facility’s software (their cluster; Data processing). A compound comprises a main feature (the primary ion with the best library match), alternative-name features (other isomeric library entries, integrated within their own retention-time windows, so their values are similar but not identical), isotopologues (^13^C, ^15^N), features 2.016 Da from the main ion in the same polarity (labeled oxidized or reduced by the software; these are separate signals, most likely metabolites that differ from the main one by two hydrogens, since such a shift rarely arises in the ion source), and adduct and neutral-loss features (Table S3). The catalog set comprises 4,533 compounds, 234 of them without their main feature in the set; its 6,978 features are 4,299 main, 1,542 alternative-name, 433 isotopologue, 282 oxidized or reduced, and 422 adduct or loss features. The analysis set comprises 949 compounds, 100 of them without their main feature; its 1,715 features are 849, 536, 90, 128 and 112 of the same kinds. Statistics were computed on every feature and counts reported per compound. Where one value per compound was needed, the compound was represented by its main feature, or by its most frequently detected feature if the main feature was not in the set; for age, it was represented by its most age-associated feature. Compounds still overcount metabolites, because the software links the ions of one metabolite across ionization modes, or with its in-source fragments, only in some cases ^37^.

### Annotation and chemical classification

Annotations are putative: MS/MS spectral library matches (Data processing) that were not confirmed with authentic standards, corresponding to level 2 of the Metabolomics Standards Initiative^47^. Compounds were assigned to chemical classes through the HMDB 5.0 chemical taxonomy, matched by InChIKey^48^, preceded by name rules for groups the taxonomy does not separate (peptides, acylcarnitines and oxylipins) and for compounds of external origin, such as drugs, pesticides and surfactants. The rules and their order are given in Supplementary Note 6. For Tables S4 and S13, compounds were grouped as one likely metabolite when their features were ion forms of one formula within 15 ppm, their retention times lay within 0.25 min, and their intensities co-varied across samples (Pearson r ≥ 0.5). This grouping joins compounds that the facility’s clustering left separate, such as the protonated and deprotonated ions of one metabolite, and rests on mass, retention time and co-variation, not on MS/MS spectra.

### Statistical analysis

#### Association with age

For each feature in the analysis set, the association with age at collection was measured by Spearman’s rank correlation across the 782 samples, with p values from the t approximation and q values by the Benjamini–Hochberg procedure over the 1,715 features^49^. Features with q < 0.05 were called associated with age. A compound counts as associated with age if any of its features is, and takes the correlation of its most strongly associated feature. The screen was repeated on the 849 main features alone, with its own false discovery rate. As a stricter count of independent signals, features were also grouped by correlation: taken in order of decreasing variance, each feature joined the group whose first member it correlated with most strongly if that absolute Pearson correlation was at least 0.85, and otherwise started a new group. The balance of falling and rising compounds was tested with a two-sided binomial test.

#### Adjusted associations

Partial Spearman correlations were computed by regressing the ranks of each feature and of age on covariates and correlating the residuals, with the degrees of freedom reduced by the number of covariates. Covariates were considered one at a time: sex; batch; recruitment group (15 conditions and controls); collection date, as a natural spline with 5 degrees of freedom; and three properties of each sample’s assay, namely the number of features detected, the mean intensity and the first principal component of the analysis matrix, alone and together (Table S14).

#### Robustness and shape

The screen was repeated under alternative normalizations and feature sets (Table S9), within age ranges and in control dogs only (Table S8). Departure from a linear trend was tested for each feature by comparing a linear fit of age with a natural spline of age (5 degrees of freedom) in a nested F test, with Benjamini–Hochberg correction.

#### Chemical classes

For each class, the share of compounds associated with age was compared with that of all other compounds by Fisher’s exact test, with Benjamini–Hochberg correction across classes. Each compound took the class of its most age-associated feature. A class score for each sample was the mean z-score of the class’s compounds; its association with age was measured by Spearman correlation overall, within age ranges and within dogs.

#### Age clock

Age was estimated from all 1,715 features by elastic-net regression^50^ (mixing parameter α = 0.5) using glmnet^51^. Accuracy was estimated by 10-fold cross-validation in which all samples of a dog fell in the same fold. Within each training set, features were centered and scaled, and the penalty was chosen by five-fold cross-validation as the value minimizing mean squared error. Out-of-fold estimates were summarized as Pearson r, R² and mean absolute error. The coefficients reported come from one fit to all samples. One variant used one feature per correlated group, and another the 500 features most correlated with age (selected within each training set). Refits within subgroups and without classes of features are in Tables S11 and S12.

#### Within-dog comparison

For each dog sampled more than once, the out-of-fold estimated ages of its first and last samples were compared. Both samples of a dog were estimated by a model trained without that dog. The share of dogs whose later sample was estimated older was tested against one half with an exact binomial (sign) test, with a Clopper–Pearson 95% confidence interval.

#### Collection date

Age and collection date are strongly correlated in this cohort, so we tested whether the association with age persists at a fixed collection date. Features and age were each adjusted for collection date (natural spline, 5 degrees of freedom), and the adjusted Spearman correlation of each feature with age was computed. The test statistic was the mean adjusted correlation over the n features most associated with age, each signed by the direction of its unadjusted association, where n is the number of features associated with age in the unadjusted screen. Its null distribution came from 2,000 permutations of age among samples within 20 bins of collection date (repeated with 10 and 40 bins), so that only samples of similar collection date were exchanged^52,53^, with the selection of the n features repeated in every permutation. If hair chemistry depended on age alone, adjustment for date would shrink each correlation by a predictable amount, set by the share v of age variance that remains after removing date: a feature with correlation ρ would have an adjusted correlation of about √(vS/(vS + 1)), where S = ρ²/(1 − ρ²). The expected statistic is the mean of this value over the n features. The age share is the observed statistic minus the mean of the null, divided by the expected statistic, so that 0 corresponds to an association entirely due to collection date and 1 to one entirely due to age; its 95% interval is ±1.96 standard deviations of the null, divided by the expected statistic.

#### Validation against recorded exposure

Owners in the cohort report flea, tick and heartworm preventives each year, including product, route and months of use. For the first sample of each dog, a product counted as in use if its record lay within one year of collection and it was used year-round or in the month of collection. On the catalog set, the feature annotated as imidacloprid was compared, as specified before testing, between dogs using a topical imidacloprid product (collar or spot-on) and dogs using no preventive that month. Its area under the receiver operating characteristic curve (AUC, the Mann–Whitney U statistic divided by the product of the group sizes) was tested with a Wilcoxon rank-sum test. To separate exposure from season, the AUC was also computed within each calendar month with at least 5 dogs in each group and averaged with weights equal to the number of dogs. Further checks compared collar and spot-on products separately, compared dogs on oral products only with dogs on none, and ranked the imidacloprid feature among 500 randomly drawn features by its distance from an AUC of 0.5. Methoprene acid and dinotefuran were tested in the same way for the products that contain them. The analysis was reproduced on the analysis set.

#### Neutering status

Neutering status at each visit was reconstructed from the recorded neutering date, because the status field of the data export describes the dog at the time of export. A dog neutered on the day of the visit was counted as neutered, and 5 samples without a neutering date were left unassigned (615 neutered, 162 intact). For each sample, an age score was the mean z-score of the 430 age-associated features, each signed by its direction of association with age, so that higher values are typical of older dogs; class scores were computed as above. Neutered and intact samples were compared by linear regression of each score on neutering status, a natural spline of age (3 degrees of freedom) and sex, within age ranges. The comparison among dogs aged 3 years or more was repeated with recruitment group and collection date (natural spline, 3 degrees of freedom) added, with the first sample of each dog only, and with a score restricted to annotated features of endogenous origin (Table S15).

#### Batch structure

For each feature, differences between the 40 batches were tested by one-way analysis of variance with Bonferroni correction. Across features, the share of total variance between batches was computed as in PERMANOVA on Euclidean distances^54^, with a p value from 1,000 permutations of the batch labels.

#### Software

The analysis pipeline ran in R 4.5.2 and 4.4.2 with glmnet 4.1.9 and SummarizedExperiment. Figures, tables and the supplementary analyses used R 4.4.2 (ggplot2 4.0.3, patchwork 1.3.2) and Python 3.10.8 (SciPy 1.15.3, pandas 2.3.3). Random seeds are fixed in the code.

## Ethics statement

Owners of all dogs in the Golden Retriever Lifetime Study gave written informed consent. The study protocol was approved by Morris Animal Foundation’s Animal Welfare Advisory Board. This study was a secondary analysis of banked samples and records provided under a data use agreement with the Morris Animal Foundation; no animals were handled for this work.

## Acknowledgements

We would like to thank Morris Animal Foundation, its staff members and all participants in the Golden Retriever Lifetime Study, including the dog owners, their golden retrievers and the Study veterinarians who made this work possible.

## Author contributions

F.C. and J.K. conceived the study, extraction protocol, designed and performed the analysis, prepared the figures and wrote the manuscript. D.J., L.F. and T.R. acquired and processed the mass spectrometry data. B.S. curated the cohort data and selected and provided the banked samples. All authors read and approved the final manuscript.

## Supplementary Figures

Figure S1. Batch effects before and after correction.

Figure S2. Age and collection date

## Supplementary Notes

Supplementary Note 1. Batch correction and batch structure

Supplementary Note 2. Annotation

Supplementary Note 3. Composition

Supplementary Note 4. Age screen

Supplementary Note 5. Compounds associated with age

Supplementary Note 6. Chemical classification rules and limitations of individual annotations

## Supplementary Tables

**Table S1. Excluded batches.** The two batches excluded at quality control, with the number of study samples, the agreement between raw intensities and the facility’s non-detection assignments, and the fold deviation in overall signal for study samples, day controls and batch controls, with the retained batches summarized for comparison.

**Table S2. Batch quality-control scoreboard.** For each of the 40 retained batches, the deviation in detection rate, the relative log-abundance offset, three measures of distance from the other batches, and the mean rank over these measures.

**Table S3. Feature catalog.** The 6,978 features of the catalog set, with facility compound identifier, row type (main ion, alternative name, isotopologue, oxidized or reduced form, adduct), library annotation, ion form, formula, observed m/z, retention time and integration window, chemical class and its source, detection rate, median raw intensity, whether the feature represents its compound, membership of the analysis set, and the association with age.

**Table S4. The 30 annotated metabolites with the highest raw signal.** Catalog set (4,533 compounds detected in at least 75% of samples). Each compound is represented by its main feature, or by its most frequently detected feature if the main feature is not in the set. Signal is the median raw peak area over the samples in which the compound was detected, before normalization; raw signal is not concentration, because ionization efficiency differs between compounds. Compounds that are ion forms of one peak (protonated and deprotonated ions, adducts, in-source fragments, ^13^C isotope) are grouped into one line, with the m/z, retention time (RT), detection and signal of the strongest member. Grouping rests on accurate mass, retention time and co-variation across samples, not on MS/MS spectra. Library names are given in the table, and the members of each group in sheet S4 members. Names are putative MS/MS library matches.

**Table S5. Imidacloprid signal by reported product.** For each owner-reported product group, the number of dogs, the median log2 signal with its interquartile range, and the number of dogs in which the signal was not detected.

**Table S6. Imidacloprid and other actives against owner-reported use.** Area under the receiver operating characteristic curve for each contrast between owner-reported preventive groups, for imidacloprid, methoprene acid and dinotefuran, with the number of dogs in each group and the p value.

**Table S7. Imidacloprid within calendar months.** For each month of collection, the number of dogs reporting a topical imidacloprid product, the number on no preventive, and the area under the curve separating them.

**Table S8. Age screen by subset of samples and by adjustment.** Features and compounds associated with age in each subset (all samples; age floors of 1, 2 and 3 years; younger and older half; controls only; osteoarthritis cases removed) and under each adjustment (sex, analysis batch, recruitment group, collection date), with the number falling and rising and the strongest absolute correlation.

**Table S9. Agreement across normalizations and feature sets.** Pairwise agreement of the per-feature age correlations between the eight combinations of batch-correction method and feature set, as Pearson and Spearman correlation and as the share of features agreeing in sign, with the number of features shared by each pair.

**Table S10. Clock coefficients.** The 167 features retained by the elastic-net age clock, with compound identifier, row type, chemical class, retention time, fitted coefficient and the feature’s own correlation with age.

**Table S11. Clock refits within subgroups.** Cross-validated accuracy when the clock is refitted within each subgroup, with the number of samples and dogs, the age range, the features retained, r, R² and mean absolute error, the same for the full model rescored on those samples, and the difference in r.

**Table S12. Clock accuracy by the features used.** Cross-validated accuracy for each restricted feature set, with the number of features offered and retained, r, R², mean absolute error, and the within-dog result.

**Table S13. The 30 annotated metabolites most strongly associated with age.** Analysis set (949 compounds). Each compound is represented by its most age-associated feature (Spearman ρ; q by the Benjamini-Hochberg procedure over all 1,715 features). Compounds that are ion forms of one peak are grouped into one line (accurate mass, retention time and co-variation, not MS/MS spectra), with the statistics of the most age-associated member. Library names and feature counts are given in the table, and the members of each group in sheet S13 members. ‘After recruitment-group adjustment’ states whether the compound remains associated with age. RT, retention time. Names are putative MS/MS library matches.

**Table S14. Assay-quality covariates.** For each candidate covariate, and for the three assay properties together, its own correlation with age, the share of variance in age remaining after adjustment, and the number of features still associated with age against the 430 of the unadjusted screen.

**Table S15. Neutering comparison.** Difference between neutered and intact dogs in the age-associated score and in the oxylipin and fatty acid class scores, in standard deviations, by sample subset and covariate adjustment, with the number of intact and neutered samples and the p value.

## References

1. Gilmore, K.M. & Greer, K.A. Why is the dog an ideal model for aging research? Exp Gerontol 71, 14–20 (2015).

2. Creevy, K.E., Akey, J.M., Kaeberlein, M., Promislow, D.E.L. & Dog Aging Project, C. An open science study of ageing in companion dogs. Nature 602, 51–57 (2022).

3. Wise, C.F., et al. Comparative Exposure Assessment Using Silicone Passive Samplers Indicates That Domestic Dogs Are Sentinels To Support Human Health Research. Environ Sci Technol 54, 7409–7419 (2020).

4. Diaz, S.F., Torres, S.M.F., Dunstan, R.W. & Lekcharoensuk, C. An analysis of canine hair re-growth after clipping for a surgical procedure. Vet Dermatol 15, 25–30 (2004).

5. Stalder, T. & Kirschbaum, C. Analysis of cortisol in hair--state of the art and future directions. Brain Behav Immun 26, 1019–1029 (2012).

6. Bennett, A. & Hayssen, V. Measuring cortisol in hair and saliva from dogs: coat color and pigment differences. Domest Anim Endocrinol 39, 171–180 (2010).

7. Bryan, H.M., Adams, A.G., Invik, R.M., Wynne-Edwards, K.E. & Smits, J.E.G. Hair as a meaningful measure of baseline cortisol levels over time in dogs. J Am Assoc Lab Anim Sci 52, 189–196 (2013).

8. Huthsteiner, K., Finke, J.B., Peters, E.M.J., Klucken, T. & Stalder, T. Hair cortisol and other hair analytes decline with storage over a one-year period: A systematic, within-subject investigation. Psychoneuroendocrinology 181, 107593 (2025).

9. Jang, W.-J., et al. Hair Metabolomics in Animal Studies and Clinical Settings. Molecules 24(2019).

10. Sulek, K., et al. Hair metabolomics: identification of fetal compromise provides proof of concept for biomarker discovery. Theranostics 4, 953–959 (2014).

11. Chen, Y., Guo, J., Xing, S., Yu, H. & Huan, T. Global-Scale Metabolomic Profiling of Human Hair for Simultaneous Monitoring of Endogenous Metabolome, Short- and Long-Term Exposome. Front Chem 9, 674265 (2021).

12. van de Lavoir, M., et al. Untargeted hair lipidomics: comprehensive evaluation of the hair-specific lipid signature and considerations for retrospective analysis. Anal Bioanal Chem 415, 5589–5604 (2023).

13. Khandelwal, P., et al. 1H NMR-based lipidomics of rodent fur: species-specific lipid profiles and SCD1 inhibitor-related dermal toxicity. J Lipid Res 55, 1366–1374 (2014).

14. Makowska, K., et al. Hair Sample Analysis as a Method of Monitoring Exposure to Bisphenol A in Dogs. Int J Environ Res Public Health 19(2022).

15. Bawazeer, S., Watson, D.G. & Knottenbelt, C. Determination of nicotine exposure in dogs subjected to passive smoking using methanol extraction of hair followed by hydrophilic interaction chromatography in combination with Fourier transform mass spectrometry. Talanta 88, 408–411 (2012).

16. Yavuz, O., et al. Monitoring of environmental persistent organic pollutants in hair samples of cats and dogs. Sci Total Environ 933, 173020 (2024).

17. Makowska, K., et al. Assessment of exposure to perfluoroalkyl substances (PFASs) in dogs by fur analysis. Environ Pollut 286, 117435 (2021).

18. Seo, H.-J., et al. Target and non-target analytical method for potential hazardous substances in livestock and pet hair using liquid- and gas chromatography quadrupole time-of-flight mass spectrometry. J Chromatogr A 1705, 464170 (2023).

19. Magalhães-Junior, J.T., et al. Identification of biomarkers in the hair of dogs: new diagnostic possibilities in the study and control of visceral leishmaniasis. Anal Bioanal Chem 406, 6691–6700 (2014).

20. Suschinel, R., et al. Identification of putative volatile biomarkers of canine leishmaniasis in dog’s breath and hair employing a novel algorithm for automated chromatographic peak detection and matching. Anal Bioanal Chem 417, 771–783 (2025).

21. Prescott, J., et al. Rationale and design of the Dog Aging Project precision cohort: a multi-omic resource for longitudinal research in geroscience. Geroscience 47, 5725–5748 (2025).

22. Guy, M.K., et al. The Golden Retriever Lifetime Study: establishing an observational cohort study with translational relevance for human health. Philos Trans R Soc Lond B Biol Sci 370(2015).

23. Labadie, J., et al. Cohort profile: The Golden Retriever Lifetime Study (GRLS). PLoS One 17, e0269425 (2022).

24. Rawlings, A.V. & Harding, C.R. Moisturization and skin barrier function. Dermatol Ther 17 Suppl 1, 43–48 (2004).

25. Olivry, T., Paps, J.S. & Amalric, N. Transient and reversible reduction of stratum corneum filaggrin degradation products after allergen challenge in experimentally mite-sensitised atopic dogs. Vet Dermatol 33, 62–e20 (2022).

26. Rogers, G.E., Harding, H.W. & Llewellyn-Smith, I.J. The origin of citrulline-containing proteins in the hair follicle and the chemical nature of trichohyalin, an intracellular precursor. Biochim Biophys Acta 495, 159–175 (1977).

27. Méchin, M.-C. & Simon, M. Deimination in epidermal barrier and hair formation. Philos Trans R Soc Lond B Biol Sci 378, 20220245 (2023).

28. Bannasch, D., et al. Mutations in the SLC2A9 gene cause hyperuricosuria and hyperuricemia in the dog. PLoS Genet 4, e1000246 (2008).

29. Wooten, K.J. & Smith, P.N. Canine toys and training devices as sources of exposure to phthalates and bisphenol A: quantitation of chemicals in leachate and in vitro screening for endocrine activity. Chemosphere 93, 2245–2253 (2013).

30. Harvey, M.H., McMillan, M., Morgan, M.R. & Chan, H.W. Solanidine is present in sera of healthy individuals and in amounts dependent on their dietary potato consumption. Hum Toxicol 4, 187– 194 (1985).

31. Craig, M.S., Gupta, R.C., Candery, T.D. & Britton, D.A. Human exposure to imidacloprid from dogs treated with advantage(r). Toxicol Mech Methods 15, 287–291 (2005).

32. Chopade, H., et al. Skin distribution of imidacloprid by microautoradiography after topical administration to beagle dogs. Vet Ther 11, E1–10 (2010).

33. Buisson, M., et al. Antiparasitic Collars: Concentration Levels of Imidacloprid and Flumethrin in Dog Fur Suggest Low Toxicity Risks for Adult Humans. J Vet Pharmacol Ther 48, 250–259 (2025).

34. Diepens, N.J., Belgers, D., Buijse, L. & Roessink, I. Pet dogs transfer veterinary medicines to the environment. Sci Total Environ 858, 159550 (2023).

35. El Khoury, L.Y., et al. Systematic underestimation of the epigenetic clock and age acceleration in older subjects. Genome Biol 20, 283 (2019).

36. Agrawal, K., et al. Effects of atopic dermatitis and gender on sebum lipid mediator and fatty acid profiles. Prostaglandins Leukot Essent Fatty Acids 134, 7–16 (2018).

37. Mahieu, N.G. & Patti, G.J. Systems-Level Annotation of a Metabolomics Data Set Reduces 25 000 Features to Fewer than 1000 Unique Metabolites. Anal Chem 89, 10397–10406 (2017).

38. Alestas, T., Ganceviciene, R., Fimmel, S., Müller-Decker, K. & Zouboulis, C.C. Enzymes involved in the biosynthesis of leukotriene B4 and prostaglandin E2 are active in sebaceous glands. J Mol Med (Berl) 84, 75–87 (2006).

39. Pagac, M.P., et al. Life stage impact on the human skin ecosystem: lipids and the microbial community. NPJ Biofilms Microbiomes 11, 13 (2025).

40. Lees, H.J., Swann, J.R., Wilson, I.D., Nicholson, J.K. & Holmes, E. Hippurate: the natural history of a mammalian-microbial cometabolite. J Proteome Res 12, 1527–1546 (2013).

41. Wikoff, W.R., et al. Metabolomics analysis reveals large effects of gut microflora on mammalian blood metabolites. Proc Natl Acad Sci U S A 106, 3698–3703 (2009).

42. Wang, X., et al. Target-Decoy-Based False Discovery Rate Estimation for Large-Scale Metabolite Identification. J Proteome Res 17, 2328–2334 (2018).

43. Broadhurst, D., et al. Guidelines and considerations for the use of system suitability and quality control samples in mass spectrometry assays applied in untargeted clinical metabolomic studies. Metabolomics 14, 72 (2018).

44. Dunn, W.B., et al. Procedures for large-scale metabolic profiling of serum and plasma using gas chromatography and liquid chromatography coupled to mass spectrometry. Nat Protoc 6, 1060– 1083 (2011).

45. Wehrens, R., et al. Improved batch correction in untargeted MS-based metabolomics. Metabolomics 12, 88 (2016).

46. Johnson, W.E., Li, C. & Rabinovic, A. Adjusting batch effects in microarray expression data using empirical Bayes methods. Biostatistics 8, 118–127 (2007).

47. Sumner, L.W., et al. Proposed minimum reporting standards for chemical analysis Chemical Analysis Working Group (CAWG) Metabolomics Standards Initiative (MSI). Metabolomics 3, 211– 221 (2007).

48. Wishart, D.S., et al. HMDB 5.0: the Human Metabolome Database for 2022. Nucleic Acids Res 50, D622–D631 (2022).

49. Benjamini, Y. & Hochberg, Y. Controlling the False Discovery Rate: A Practical and Powerful Approach to Multiple Testing. Journal of the Royal Statistical Society Series B: Statistical Methodology 57, 289–300 (1995).

50. Zou, H. & Hastie, T. Regularization and Variable Selection Via the Elastic Net. Journal of the Royal Statistical Society Series B: Statistical Methodology 67, 301–320 (2005).

51. Friedman, J., Hastie, T. & Tibshirani, R. Regularization Paths for Generalized Linear Models via Coordinate Descent. J Stat Softw 33, 1–22 (2010).

52. Winkler, A.M., Ridgway, G.R., Webster, M.A., Smith, S.M. & Nichols, T.E. Permutation inference for the general linear model. Neuroimage 92, 381–397 (2014).

53. Winkler, A.M., Webster, M.A., Vidaurre, D., Nichols, T.E. & Smith, S.M. Multi-level block permutation. Neuroimage 123, 253–268 (2015).

54. Anderson, M.J. A new method for non-parametric multivariate analysis of variance. Austral Ecology 26, 32–46 (2001).

