## Supplementary Notes 1-6 for "The canine hair metabolome encodes age and coat exposure in over 700 dogs"

#### Supplementary Note 1. Batch correction and batch structure

On 4,000 randomly chosen features of the full data set that the correction could reach, variance between batches fell by 39.6% in samples not used to fit the correction and by 44.8% in the samples used, and the two halves agreed with r 0.887. On the 1,358 such features of the analysis set, the reduction was 47.6% (55.6% in the samples used). On detected values only, the median share of variance between batches was 0.286 without correction, 0.262 with the facility's normalization and 0.188 with ours (Figure S1B). In the uncorrected data, the compound with the highest signal was about 85-fold (6.4 log2 units) higher in batches 17 to 40 than in batches 1 to 16 (mean of batch means); the standard deviation of its batch means was 3.23 log2 units before correction and 0.76 after (Figure S1A).

After correction, 256 features still had a batch mean more than 8-fold (3 log2 units) from the average of all batches, against 633 before, and 159 of the 256 were features the correction could not reach. Conversely, 302 of the 68,600 batch means, in 188 features (143 of them out of reach), ended more than 4-fold from the average and more than 2-fold further from it than before. The median over features of each batch's level varied across batches with a standard deviation of 0.27 log2 units without correction, 0.29 with the facility's normalization and 0.22 after correction (Figure S1C).

Between consecutive batches, the largest change came between batches 16 and 17 (337 of the 1,715 features changed more than 4-fold, in both directions), followed by batches 29 and 30 (247 features; one excluded batch was acquired between them) and batches 39 and 40 (216), against a median of 25 at the other boundaries. Of the 275 features with their apex within 0.02 min of a window edge, 166 are main ions and 240 lie at the late edge. Such features are more frequent near the edge than elsewhere (about 90 features per 0.01 min within 0.02 min of an edge, against about 30 at 0.03–0.05 min). After correction, 29% of them still have a batch more than 8-fold off, against 12% of other features, and the median standard deviation of their batch means is 0.73 log2 units, against 0.55. Ornithine (Table S4) has its apex at 10.98 min in a window ending at 10.99 min; its signal falls by up to about 1,200-fold over batches 17 to 29 and recovers at batch 30. Separately, the main ion of vaccenic acid was about 260-fold low in the last batch while its $^{13}C$ isotope was normal, an integration error that no batch correction can fix.

#### Supplementary Note 2. Annotation

All compound names reported in this study are putative. Annotations are MS/MS spectral library matches not confirmed with authentic standards, and 732 of the 2,343 annotated compounds (31%) match more than one library entry. One early-eluting lipid compound at 3.15 min matches 34 entries.

Each library entry is integrated within its own retention-time window, and where neighboring windows overlap a common metabolite can be reported under an implausible name. The two entries at 6.2–6.4 min that carry the protonated ion of urocanic acid are butylphosphonic acid and methyl 4-pyrimidine carboxylate. Two benzylamine-type names are a fragment of phenylalanine, O-acetylserine is pyroglutamic acid, isonicotinic acid integrates the edge of the nicotinic acid peak, and the mass of embelin fits a C14 alkyl sulfate better than embelin itself (1 against 10 ppm). The compound annotated as allopurinol, detected in 99.7% of samples, is almost certainly its isomer hypoxanthine (Table S4; Supplementary Note 6).

The compound with the highest signal of all has the molecular formula of urocanic acid ($C_6H_6N_2O_2$) and carries the same ion forms, but elutes at 9.05 rather than 6.3 min and co-varies with urocanic acid only moderately ($r = 0.44$). Its identity remains unresolved; one candidate is cis-urocanic acid, which forms from the trans isomer in the stratum corneum under ultraviolet light[1]. Because its signal steps between batches 16 and 17 (Supplementary Note 1), its rank by signal is approximate.

The converse also occurs, with a single metabolite reported as several compounds. The facility's software links the ions of one metabolite across ionization modes, or with its in-source fragments, only in some cases[2]. Grouping entries by mass, retention time and covariation (Methods) reduced the 53 compound entries among the 30 annotated metabolites with the highest signal to 30 likely metabolites, with urocanic acid alone accounting for five of them (Table S4).

Among the 62 compounds named as oxylipins, 13 are synthetic prostaglandin analogues or conjugates, and library entries for isotope-labeled standards and for controlled substances also match detected signals (Supplementary Note 6). Leucine and isoleucine are not chromatographically resolved. For these reasons we base our conclusions on chemical classes and on groups of related compounds rather than on individual library names.

#### Supplementary Note 3. Composition

No feature was detected in every sample; 8.8% of values in the catalog set and 3.6% in the analysis set were below the detection limit. In the catalog set, 1,655 compounds were detected in at least 95% of samples; in the analysis set, the median compound was detected in 98.6% of samples and 674 compounds in at least 95%. Median signal spans about 4.4 orders of magnitude across the catalog set, and annotated compounds carry 67% of the summed signal.

By chemical class (Figure 2C), the annotated compounds comprise peptides (542 compounds, 23% of annotated compounds, 3.6% of signal), amino acids and their derivatives (316, 13%; 16.8% of signal), heterocyclic compounds (253, 11%; 10.1% of signal, largely through urocanic acid and the unidentified compound of the same formula), small oxygen- and nitrogen-containing compounds such as sugars and amines (160, 7%), benzenoids (154, 7%), lipids other than fatty acyls (94, 4%), purines, pyrimidines and nucleosides (90, 4%; 3.9% of signal), fatty acids (68, 3%; 4.3% of signal), other organic acids (63, 3%) and oxylipins (62, 3%); no remaining class accounts for more than 7% of annotated compounds. Smaller groups include 46 plant-type secondary metabolites, 31 drugs, 30 pesticides, surfactants and industrial chemicals, 10 acylcarnitines and 9 isotope-labeled library standards, and 408 annotated compounds have no match in HMDB.

The annotated compounds with the largest signals are dominated by constituents of the skin surface and the hair shaft (Table S4): urocanic acid, pyroglutamic acid, phenylalanine, leucine and isoleucine, glutamic acid, ornithine, citrulline, 4-guanidinobutanoic acid, β-alanine or sarcosine, uric acid, allantoic acid and adenosine. In human stratum corneum the natural moisturizing factor is about 40% free amino acids, 12% pyroglutamic acid and 12% lactate[3], and derives in part from the breakdown of filaggrin; free serine and lactate were both detected here, at lower signal than pyroglutamic acid. Urocanic and pyroglutamic acids decline in canine skin after allergen challenge[4], and urocanic acid absorbs ultraviolet light[5]. Citrulline reflects the conversion of arginine residues in trichohyalin during hair formation[6,7]. Allantoic acid is a hydrolysis product of allantoin, which dogs other than Dalmatians form when breaking purines down past uric acid[8]. The short peptides, among them His-Pro, are consistent with keratin breakdown, while two apparent high-signal peptides, Thr-Gly and Gly-Phe-Lys, are in fact ions of citrulline (Table S4). The lipids detected include vaccenic, hexadecenoic and docosatetraenoic acids, sphingoid bases and oxidized fatty acids, consistent with sebum.

Four of the 30 annotated metabolites with the largest signals are sulfate surfactants: myristyl sulfate, a second C14 alkyl sulfate reported under the name embelin, and the laureth sulfates ethylene- and diethylene-glycol dodecyl ether sulfate. Triethylene-glycol dodecyl ether sulfate is also present, and Surfynol PSA-336, an acetylenic-diol surfactant, ranks 29th. The phthalates detected are dimethyl, diethyl, di-n-pentyl, diisooctyl and diisodecyl phthalate, and bisphenol A has been reported to leach from dog toys and training devices[9]. Further exogenous compounds, at lower signal, include methylparaben, the pesticides imidacloprid, dinotefuran, methoprene acid, atrazine and cycloate, salicylates, and putative gingerols and shogaol. Methoprene acid is combined with fipronil in some flea and tick products, and fipronil itself is not present in the facility's library. Solanidine, a potato glycoalkaloid, was detected at high signal in 95% of samples; its serum concentration tracks potato consumption in people[10].

#### Supplementary Note 4. Age screen

For 117 of the 155 age-associated compounds the main feature was itself associated with age; for the other 38, only an alternative-name, isotope, oxidized or reduced, or adduct feature was. The number of features detected in each sample, its mean intensity and its first principal component were themselves unrelated to age ($|r| < 0.06$); adjusted for them, 416, 521 and 512 features were associated with age. Without the 275 features at the edge of their integration window, 405 features in 149 compounds were associated with age, and the 10 compounds that dropped out had $|\rho| \leq 0.17$. A nested F test comparing a linear fit of age with a natural spline of age (5 degrees of freedom) flagged 364 features as departing from linearity, but none once the 23 samples from dogs under 1 year were excluded; the apparent curvature reflects the leverage of a few young samples rather than a bend in the age trajectory.

Clock refits (Tables S11 and S12): without the dogs recruited for osteoarthritis, r 0.623 against 0.639 for the full model on the same samples; within adults aged 2–10 years, 0.478 against 0.551; without the oxylipins (209 features), 0.657; on main features only, 0.654; without the features at the edge of their integration window, 0.658. Without the features of external origin, the mean absolute error was 1.99 years.

**Collection-date sensitivity analysis.** Age and collection date were strongly correlated ($r = 0.954$), leaving only 8.1% of age variation independent of date. After date adjustment, only 2 of the 155 age-associated compounds (3 of 430 features) remained significant. In the date-restricted permutation test, the mean signed, date-adjusted correlation was +0.040, compared with +0.008 under the null ($P = 0.021$; Figure S2B). The estimated share attributable to age was 0.83 (95% interval 0.07–1.59). Results were similar using one feature per correlated group (share 0.77; $P = 0.009$), varying the number of date bins (shares 0.70–0.83), and excluding features near integration-window edges (shares 0.75–0.88; $P = 0.017$–$0.026$). Results were weaker using main features only (shares 0.47–0.62; $P = 0.022$–$0.078$; Figure S2C).

#### Supplementary Note 5. Compounds associated with age

Of the 79 annotated age-associated compounds, 55 fall and 24 rise. The 30 in Table S13 form likely metabolites (23 falling and 7 rising; 13 remain associated after adjustment for recruitment group) and are tagged by likely origin (19 endogenous, 6 from diet or gut microbes, 1 not dog biology, 4 with implausible names).

The peak at m/z 335.2228 could also be a DiHETE or a water-loss ion of a prostaglandin. In 11 of the 14 age-associated oxylipin compounds, the main feature is itself associated; 7 of those main features carry an oxylipin name and 4 are classed otherwise (isosteviol, a conjugated linoleic acid and two names without an HMDB class), and the other 3 compounds are associated only through alternative-name or oxidized or reduced features. 8-HETE, 15-HETE and LTB3 are one HETE peak, and 15-HEPE and an EpETE are one peak; after 9,10-DiHOME come 9-HpODE and the HEPE/EpETE peak. Without features at the edge of their integration window, 9 of the 10 peaks remain associated, and lipoxin A4 drops out. Adjustment for Surfynol PSA-336 gave similar results for the other oxylipins, as did adjustment for the total signal of the window.

The 4 age-associated carnitine features (of 8) all rise. The rising sphingoid bases are heptadecasphing-4-enine (+0.14) and a C15 sphingosine (+0.13); nicotinic acid rises by +0.13 (the 3-pyridinecarboxaldehyde entry is the same peak, r 0.93), 2-aminoadipic acid by +0.12, and a few tripeptides, among them Tyr-Val-Lys (+0.13). Other falling compounds of diet or gut-microbial origin are 3-aminobenzoic acid (−0.17) and 4-hydroxybenzenesulfonic acid (−0.12), which has the formula of phenyl sulfate. A compound annotated as dimethylglycine (+0.13) is probably not dimethylglycine: it elutes with the lipids (3.29 min) and co-varies with heptadecasphing-4-enine (r 0.84). A compound annotated as p-acetaminobenzoic acid is the same signal as hippuric acid (r 0.89), and 4-hydroxybenzenesulfonic acid co-varies with indoxyl sulfate (r 0.57).

##### Supplementary Note 6. Chemical classification rules and limitations of individual annotations

Every row of the facility table is assigned a single class, and a compound takes the class of its representative row (Methods). The rules are implemented in analysis/classify_features.py, the taxonomy lookup in analysis/hmdb_taxonomy.py (HMDB 2021, 217,899 records) and the counts in analysis/composition_and_age_by_class.py.

Classification rules (applied in order, with the first match assigning the class)

| Order | Class | Rule |
|---|---|---|
| 1 | Unannotated | No library match (facility placeholder name Feature_<polarity>_<m/z>_<RT>) |
| 2 | Isotope-labelled (library standards) | Name carries a label: -d4, d3-, 13C, 15N, "internal standard" |
| 3 | Exogenous: drugs | Name matches a curated drug list (e.g. dapsone, azatadine, allopurinol, salicylates, anabolic steroids, opioids, synthetic cannabinoids) |
| 4 | Exogenous: pesticides, surfactants, industrial | Name matches a curated list (phthalates, bisphenols, parabens, siloxanes, alkyl and alkyl-ether sulfates, Surfynol, imidacloprid, dinotefuran, methoprene, atrazine, cycloate, laboratory buffers) |
| 5 | Peptides | Name is a sequence of two or more amino-acid three-letter codes |
| 6 | Acylcarnitines | Name contains "carnitine" |
| 7 | Oxylipins | Name matches eicosanoid / octadecanoid nomenclature (prostaglandins, leukotrienes, lipoxins, HETE, HEPE, HODE, HpODE, EpOME, DiHOME, isoprostanes) |

| | | |
|---|---|---|
| 8 | HMDB class | InChIKey matched to HMDB (full key, else first block), mapped to: amino acids and derivatives; peptides; oxylipins (lipid with an eicosanoid/octadecanoid parent **and** an oxidation mark in the name); acylcarnitines; fatty acids and other fatty acyls; other lipids; purines, pyrimidines and nucleosides; other organic acids; heterocyclics; benzenoids; small organic O/N compounds; plant secondary metabolites; other |
| 9 | Annotated, unclassified | Library name present but no rule or HMDB match |

The oxidation-mark requirement in rule 8 is necessary because HMDB files unoxidized linoleic and linolenic acids under the same parent classes as their oxidation products.

Limitations of individual annotations

Annotations are MS/MS spectral library matches, not confirmed with authentic standards. The cases documented below should be taken into account whenever an individual compound name is interpreted.

**Synthetic prostaglandin analogues and conjugates among oxylipin names (13 of 62 oxylipin compounds in the 4,533-compound set):** misoprostol (free acid); 15-cyclohexyl pentanor PGF2α; 16,16-dimethyl-6-keto-PGE1; 8-iso-16-cyclohexyl-tetranor-PGE2; 16-phenyl tetranor PGF2α; PGA2 methyl ester; 9,11-methane-epoxy PGF1α; 17-phenyl trinor PGA2 (two entries); 15(S)-HETE-biotin; PGF2α diethyl amide; 16,16-dimethyl-PGA2; 16,16-dimethyl PGF2α.

Library names that correspond to alternative ions of common metabolites (grouped in Tables S4 and S13; members listed in sheets S4 members and S13 members):

Each library entry is integrated in its own 0.5-min retention-time window, and the windows overlap. A neighboring entry with the same m/z therefore integrates the edge of the same peak, and its reported apex sits at the edge of its window (e.g. methyl 4-pyrimidine carboxylate at 6.16 min, window ending 6.18; isonicotinic acid at 6.16, window ending 6.17; Gly-Phe-Lys at 13.34, window ending 13.35). Entries were grouped only when the m/z is an ion form of one formula (≤ 15 ppm), the retention time is within 0.25 min, and the signals co-vary across samples (Spearman r ≥ 0.5).

| Library name | Likely identity |
|---|---|
| Butylphosphonic acid; methyl 4-pyrimidine carboxylate (RT 6.16) | urocanic acid [M+H]+ |
| 4-Imidazoleacrylic acid | urocanic acid (synonym) |

| | |
|---|---|
| [4-(Aminomethyl)phenyl]methanamine; 2-amino-1-phenylethanol | phenylalanine fragment [M+H−HCOOH]+ |
| O-Acetyl-L-serine (RT 8.04) | pyroglutamic acid [M+H]+ |
| O-Acetyl-L-serine, 2-furancarboxylic acid (RT 11.81); D-pyroglutamic acid (RT 11.71) | glutamic acid water-loss ions [M+H−H2O]+, [M−H−H2O]− |
| Gly-Phe-Lys [M+2H]2+; Thr-Gly; N-acetyl-L-histidine | L-citrulline [M+H]+, [M+H−NH3]+, [M+Na]+ |
| Isonicotinic acid; 3-pyridinecarboxaldehyde | nicotinic acid [M+H]+ (edge of the same peak) |
| Embelin | m/z fits a C14 alkyl sulfate (1 ppm) better than embelin (10 ppm); co-varies with myristyl sulfate (r 0.84) at another RT |
| 3-(Methylsulfanyl)aniline | 13C isotopologue of the RT 9.05 C6H6N2O2 compound |
| p-Acetaminobenzoic acid | hippuric acid (same m/z and RT, r 0.89) |
| Leukotriene B3, 15(R)-HETE, 9-thiastearic acid (RT 2.96–2.99) | one HETE peak, negative and positive-mode ions (positional isomer not resolved) |
| Prostaglandin F2α; 11β-prostaglandin E1 (m/z 335.2228) | one peak: a DiHETE [M−H]− or a prostaglandin's water-loss ion |
| N-Methylglutamic acid | 2-aminoadipic acid (isomer, same peak) |
| Allopurinol (99.7% detection) | hypoxanthine (isomer) |

**The highest-signal compound (m/z 139.0498 [M+H]+, RT 9.05)** has urocanic acid's formula (C6H6N2O2) and the same ion forms ([M+H]+, [M−H]−, [M+H−H2O]+, 13C), but elutes as a separate peak from urocanic acid (6.3 min). One candidate is cis-urocanic acid, which is formed from trans-urocanic acid by ultraviolet light in skin.

**Annotations of doubtful plausibility among the age-associated compounds (Table S13):** 8,11-eicosadiynoic acid (synthetic fatty-acid analogue; likely an isomeric endogenous lipid); N-acetylaspartylglutamate (neuronal dipeptide; co-varies with hippuric acid, r 0.65); dimethylglycine (a polar osmolyte's name at a lipid retention time, 3.29 min; co-varies with heptadecasphing-4-enine, r 0.84); linoleic acid-d4 (a labelled standard's library entry; most likely the $^{13}C_2$ isotope of vaccenic acid). 4-Hydroxybenzenesulfonic acid has the formula of phenyl sulfate, a gut-microbial co-metabolite, and co-varies with indoxyl sulfate (r 0.57).

**Library entries for controlled substances and isotope-labelled standards that matched detected signals:** psilocybin, desomorphine, synthetic cannabinoids (JWH series, CP 55,940, AM2201) and deuterated eicosanoid standards (linoleic acid-d4, 8-iso-PGE2-d4, lipoxin A4-d5). None of these standards was added to the samples, whose only internal standards were the labeled amino acids (Methods), so these names are false matches to other signals.

**Unresolved isomers:** leucine and isoleucine are not separated; ornithine [M+H−NH3]+ is isobaric with proline [M+H]+; N-methyl-α-aminoisobutyric acid has the formula of valine.

### References


1. de Fine Olivarius, F., Wulf, H.C., Crosby, J. & Norval, M. Isomerization of urocanic acid after ultraviolet radiation is influenced by skin pigmentation. *J Photochem Photobiol B* **48**, 42–47 (1999).

2. Mahieu, N.G. & Patti, G.J. Systems-Level Annotation of a Metabolomics Data Set Reduces 25 000 Features to Fewer than 1000 Unique Metabolites. *Anal Chem* **89**, 10397–10406 (2017).

3. Rawlings, A.V. & Harding, C.R. Moisturization and skin barrier function. *Dermatol Ther* **17 Suppl 1**, 43–48 (2004).

4. Olivry, T., Paps, J.S. & Amalric, N. Transient and reversible reduction of stratum corneum filaggrin degradation products after allergen challenge in experimentally mite-sensitised atopic dogs. *Vet Dermatol* **33**, 62–e20 (2022).

5. Gibbs, N.K., Tye, J. & Norval, M. Recent advances in urocanic acid photochemistry, photobiology and photoimmunology. *Photochem Photobiol Sci* **7**, 655–667 (2008).

6. Rogers, G.E., Harding, H.W. & Llewellyn-Smith, I.J. The origin of citrulline-containing proteins in the hair follicle and the chemical nature of trichohyalin, an intracellular precursor. *Biochim Biophys Acta* **495**, 159–175 (1977).

7. Méchin, M.-C. & Simon, M. Deimination in epidermal barrier and hair formation. *Philos Trans R Soc Lond B Biol Sci* **378**, 20220245 (2023).

8. Bannasch, D.*, et al.* Mutations in the SLC2A9 gene cause hyperuricosuria and hyperuricemia in the dog. *PLoS Genet* **4**, e1000246 (2008).

9. Wooten, K.J. & Smith, P.N. Canine toys and training devices as sources of exposure to phthalates and bisphenol A: quantitation of chemicals in leachate and in vitro screening for endocrine activity. *Chemosphere* **93**, 2245–2253 (2013).

10. Harvey, M.H., McMillan, M., Morgan, M.R. & Chan, H.W. Solanidine is present in sera of healthy individuals and in amounts dependent on their dietary potato consumption. *Hum Toxicol* **4**, 187–194 (1985).
