## Supplementary Figures S1-S2 for "The canine hair metabolome encodes age and coat exposure in over 700 dogs"

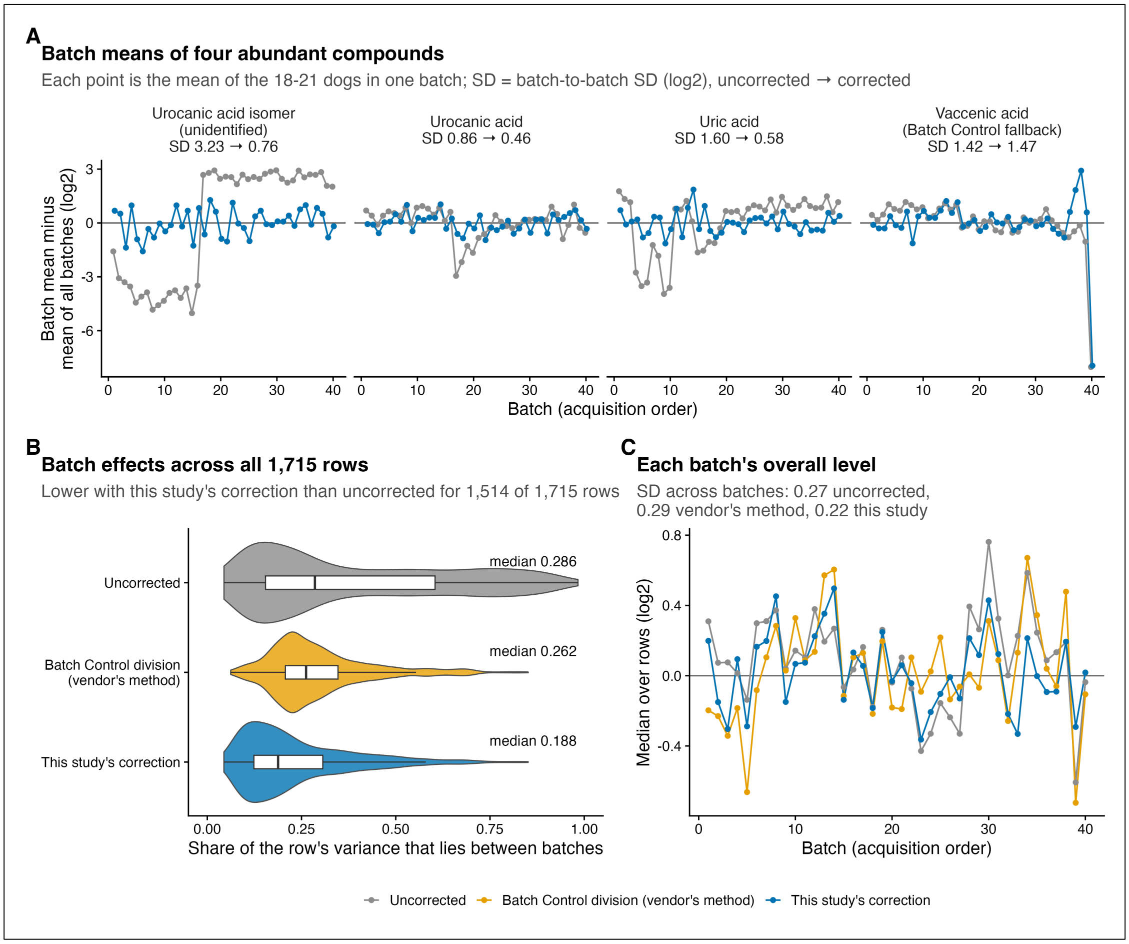


**Figure S1. Batch effects before and after correction.** (A) Batch means of four abundant compounds, without and with correction; batches are numbered in acquisition order. (B) Share of each feature's variance lying between batches without correction, with the facility's normalization to batch control samples, and with the correction used here (detected values only; medians 0.286, 0.262 and 0.188). (C) Each batch's overall level (median over features). The correction anchors each feature to the day control samples; the 357 features it cannot reach keep the facility's normalization.

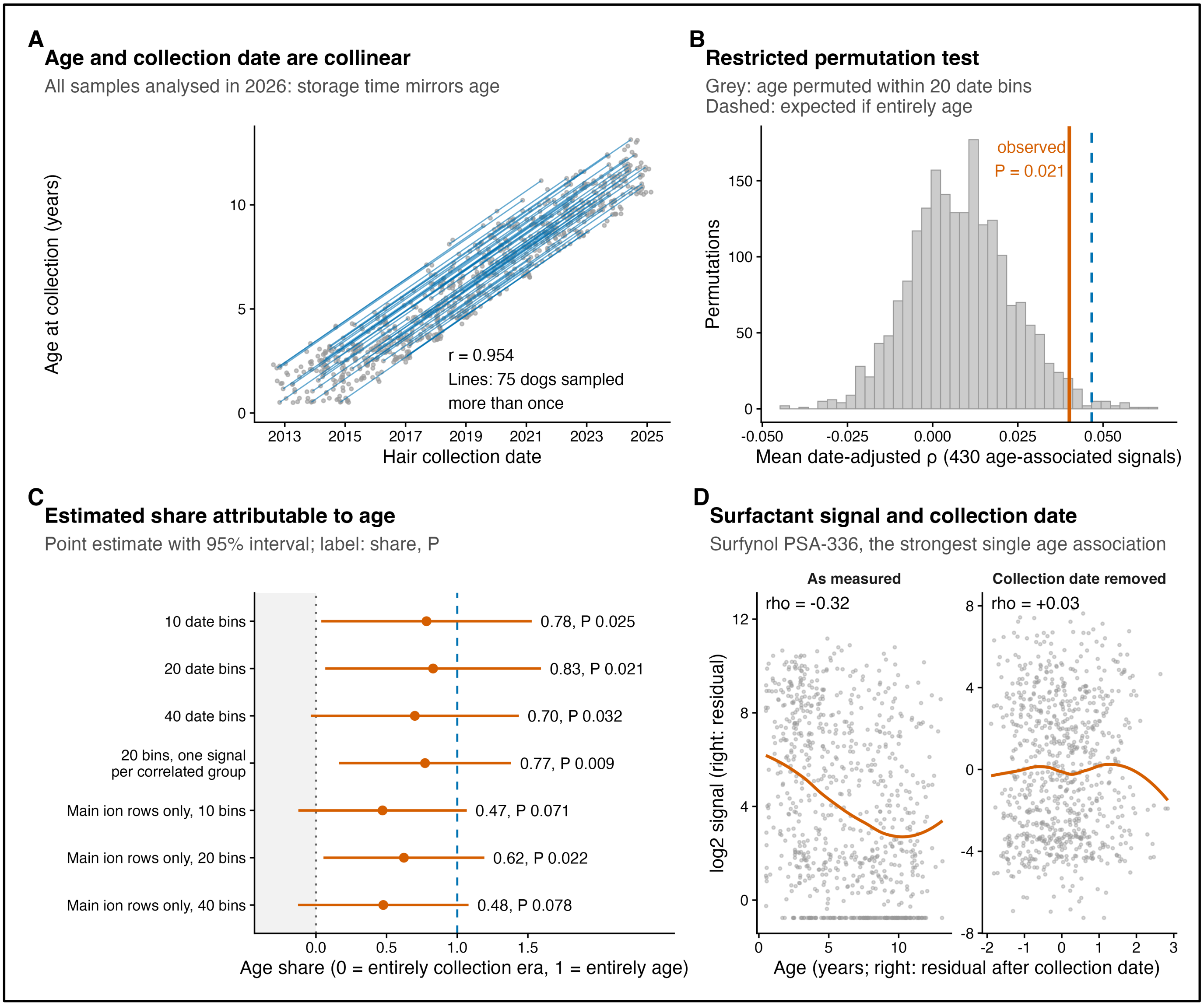


**Figure S2. Age and collection date.** (A) Age against collection date (r 0.954), with the 75 resampled dogs drawn as segments from first to last sample. (B) Null distribution of the restricted-permutation statistic (age shuffled within 20 collection-date bins), with the observed value (P = 0.021) and the value expected if the association were entirely due to age. (C) Estimated share of the signal attributable to age, with 95% intervals, for 10, 20 and 40 bins, for one feature per correlated group, and for main features only. (D) The compound annotated as Surfynol PSA-336 against age, as measured (ρ −0.32) and after removal of collection date (+0.03).
